# Polarization of Myosin II refines tissue material properties to buffer mechanical stress

**DOI:** 10.1101/241497

**Authors:** Maria Duda, Nargess Khalilgharibi, Nicolas Carpi, Anna Bove, Matthieu Piel, Guillaume Charras, Buzz Baum, Yanlan Mao

**Affiliations:** MRC Laboratory for Molecular Cell Biology, University College London, Gower Street, London WC1E 6BT, United Kingdom; London Centre for Nanotechnology, University College London, Gower Street, London WC1E 6BT, UK; Centre for Computation, Mathematics and Physics in the Life Sciences and Experimental Biology (CoMPLEX), University College London, Gower Street, London WC1E 6BT, UK; Institut Curie, PSL Research University, CNRS, UMR 144, 75005 Paris, France; Institute for the Physics of Living Systems, University College London, Gower Street, London WC1E 6BT, UK; Cell and Developmental Biology, University College London, Gower Street, London WC1E 6BT, UK

**Keywords:** tissue mechanics, Myoll polarity, force buffering, Diaphanous, elasticity, stiffness, shape maintenance

## Abstract

As tissues develop, they are subjected to a variety of mechanical forces. Some of these forces, such as those required for morphogenetic movements, are instrumental to the development and sculpting of tissues. However, mechanical forces can also lead to accumulation of substantial tensile stress, which if maintained, can result in tissue damage and impair tissue function. Despite our extensive understanding of force-guided morphogenesis, we have only a limited understanding of how tissues prevent further morphogenesis, once shape is determined after development. Buffering forces to prevent cellular changes in response to fluctuations of mechanical stress is critical during the lifetime of an adult organism. Here, through the development of a novel tissue-stretching device, we uncover a mechanosensitive pathway that regulates tissue responses to mechanical stress through the polarization of Myosin II across the tissue. Mechanistically, this process is independent of conserved Rho-kinase signaling but is mediated by force-induced linear actin polymerization and depolymerization via the formin Diaphanous and actin severing protein Cofilin, respectively. Importantly, these stretch-induced actomyosin cables stiffen the tissue to limit changes in cell shape and to protect the tissue from further mechanical damage prior to stress dissipation. This tissue rigidification prevents fractures in the tissue from propagating by confining the damage locally to the injured cells. Overall this mechanism of force-induced changes in tissue mechanical properties provides a general model of force buffering that rapidly protects tissues from physical damage to preserve tissue shape.

## Introduction

Tissue shape and function are tightly coupled: simple changes in cell geometry affect fundamental processes such as cell growth, death, or the direction of cell divisions (Chen et al., 1997; Thery et al., 2007). During development, force-guided tissue shape changes are both critical and instructive for morphogenesis (Heisenberg and Bellaiche, 2013; Legoff et al., 2013; Munjal et al., 2015; Vuong-Brender et al., 2017). In developed and differentiated tissues such as those of the adult heart or lung, it is important to preserve the correct shape for normal organ function (Liu et al., 1999; Lyon et al., 2015), which is especially challenging in face of the continuous fluctuations in mechanical forces experienced by the tissues. Mechanical forces can originate from tissue intrinsic processes such as cell division and cell death or from the external environment through processes like extracellular matrix (ECM) remodeling, mechanical changes in neighbouring tissues, gravity, fluid flow or air flow (Etournay et al., 2015; Haigo and Bilder, 2011; Jufri et al., 2015; Legoff et al., 2013; Liu et al., 1999; Mao et al., 2013; Monier et al., 2015; Porazinski et al., 2015). Unremitted mechanical stress can impact tissue integrity, fidelity of cell division and cause tissue fracture (Casares et al., 2015; Lancaster et al., 2013). Although tissues have developed a plethora of active cellular-scale mechanisms to dissipate mechanical stresses, such as cell extrusions, divisions, transitions and fusions, the full impact of these active cellular behaviors can take up to several hours (Campinho et al., 2013; Etournay et al., 2015; Heisenberg and Bellaiche, 2013; Marinari et al., 2012; Wyatt et al., 2016; Wyatt et al., 2015). As such, on the short time scale, tissues must also have an immediate protective response mechanism. They can adapt their material properties to protect their morphology while the forces dissipate; for instance, in epithelia, such ability to resist stress could be linked to their mechanical properties such as elasticity or stiffness (Bruckner and Janshoff, 2015; Skoglund et al., 2008; Zhou et al., 2009). Here we utilize the *Drosophila* wing imaginal disc to investigate the molecular and cellular basis of epithelial mechanics and the role of its dynamic remodeling in tissue shape maintenance and injury responses in stretch-challenged tissues.

## Results

### MyoII is essential for setting tissue stiffness and elasticity

Cell shape is defined by the balance of forces exerted on cells through the external environment (such as cell-cell and cell-ECM adhesion) and the forces exerted by intracellular cell components such as the actomyosin cortex (Mao and Baum, 2015). Therefore, the pathways controlling these processes are likely to be critical in responses to mechanical stress. We focused on the non-muscle Myosin II (MyoII) contractility pathway, as MyoII had been shown to be recruited to the cell cortex in force-driven morphogenetic processes such as mesoderm invagination in gastrulation as well as by deformation applied through micropipette aspiration (Fernandez-Gonzalez et al., 2009; Pouille et al., 2009). MyoII anisotropy has also been correlated with emergent tension patterns in the wing disc epithelium (Legoff et al., 2013; Mao et al., 2013). Although studies of these processes suggested that MyoII could be sensitive to mechanical stimuli, it is unclear whether MyoII accumulation is the cause or the consequence of increased tension. To test this directly, we looked at the function of MyoII in mechanically challenged tissue. In order to directly apply a controllable and quantifiable mechanical stress to a tissue, we designed a novel tissue stretching/compression device (Fig. 1A-D). Contrary to previous set ups that rely on adhesion of cells to PDMS, this device uses a novel mechanism to clamp tissue explants in order to stretch or compress stiff tissues, whilst suspended in growth media (Aragona et al., 2013; Aw et al., 2016; Eisenhoffer et al., 2012). The wing disc is positioned over the microchannel and while the sides of the wing disc are clamped by the two PDMS layers, the central portion of the tissue is effectively submerged in the microchannel, which is perfused with *ex vivo* culture media (Mao et al., 2013; Mao et al., 2011). Stretching of the PDMS sandwich concomitantly stretches the wing disc inside the microchannel. Such setup eliminates unspecific effects of PDMS interactions such as shear forces or chemical signaling, which could not be excluded in previously published devices.

**Figure 1:**
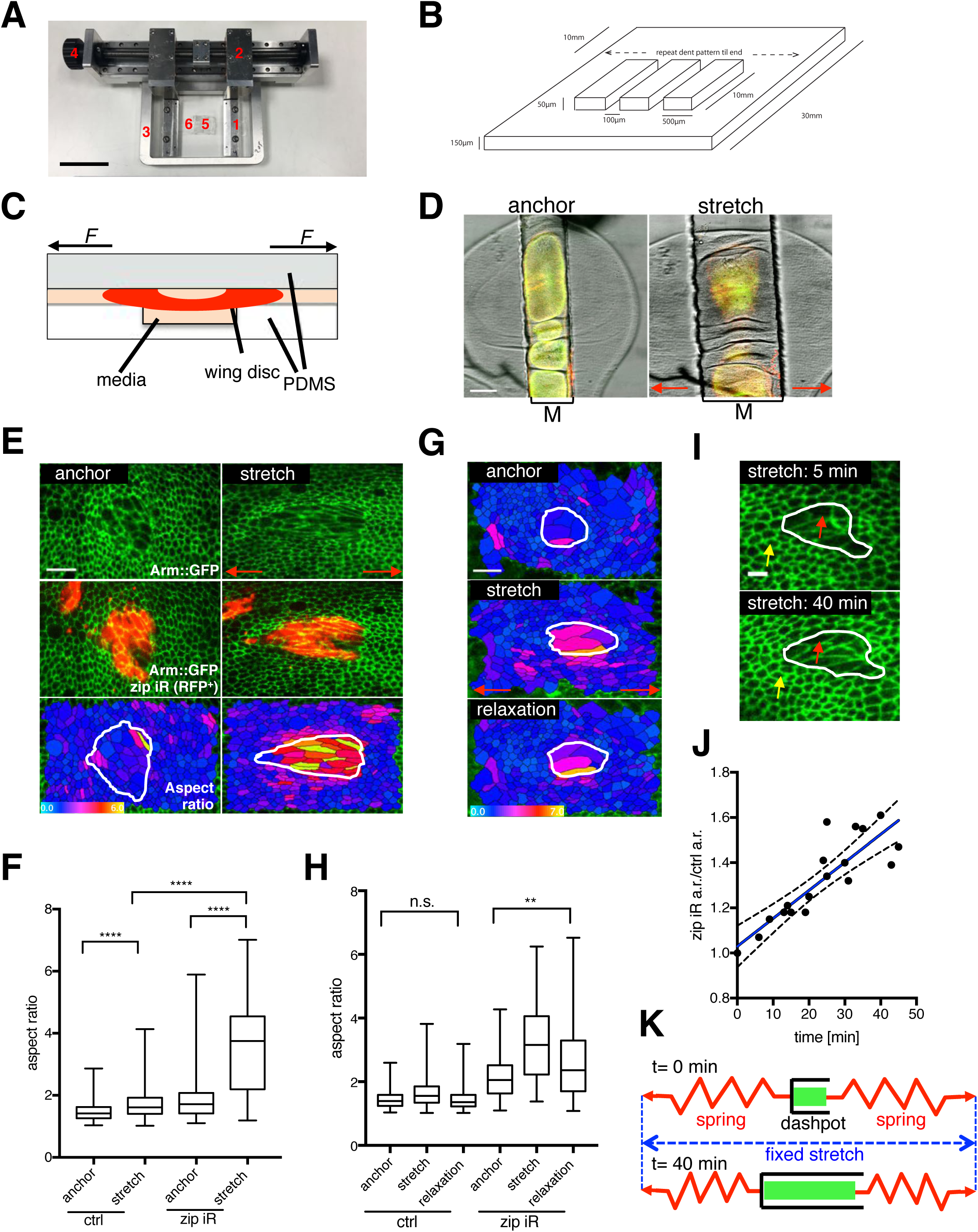
Myosin RNAi clones are softer and less elastic. (A) Stretching/compression device; 1: clamping mechanism, 2: arms, 3: stage insert, 4: drive mechanism, 5: media-filled Polydimethylsiloxane (PDMS) chamber, 6: two layers of stretchable elastomer (PDMS), one of which is pre-patterned with microchannels. (B) Scheme of PDMS prepatterning; the dimension of microchannels are 80–120 *μ*m in width and 50 *μ*m in depth. (C) Cross-sectional schematic view of the stretching device; wing disc (in red) is positioned over the microchannel with sides clamped by the two PDMS layers. The central portion of the tissue is submerged in the microchannel, perfused with *ex vivo* culture media (Mao et al., 2013; Mao et al., 2011). (D) Stretching of the PDMS sandwich concomitantly stretches the wing disc inside the microchannel (top-down view); wing disc resting on a stretching device (anchor) and 10 min after stretch; M=microchannel. (E) *zipper* RNAi clones *(zip* iR, in red) in Arm::GFP expressing anchored and stretched third instar wing disc; lower panel shows color coded cell aspect ratio in anchored and stretched discs (bright colors indicate high elongation); *zip* RNAi clone is outlined in white. (F) Box plot showing distribution of cell aspect ratio for control (wild-type) and *zipper* RNAi cells in anchored and stretched (20 min) wing disc; median represented by horizontal line, 75^th^ and 25^th^ centiles are represented by top and bottom of the boxes respectively; n=8 wing discs. (G) Color-coded aspect ratio in control and *zip* RNAi cells *(zip* RNAi clone outlined in white) subjected to anchoring, stretching (20 min) and relaxation (10 min, post stretch). (H) Box plot showing distribution of cell aspect ratio in anchored, stretched (20 min) and relaxed (10 min) wing disc for control (wild-type) and *zipper* RNAi cells; median represented by horizontal line; 75^th^ and 25^th^ centiles are represented by top and bottom of the boxes respectively, n=3 wing discs. (I) *zip-RNAi* clone in Arm::GFP wing disc at the beginning (5 min) and at the end (40 min) of stretching experiment. Yellow and red arrows indicate the control and *zip* RNAi cells respectively. (J) Change of *zip-RNAi* aspect ratio (a.r.) relative to change in control cells aspect ratio (a.r.) in the course of stretch, blue line shows fitted regression; dotted line represents 95% confidence bounds of the best fit line; n=3 wing discs. (K) Schematics describing behavior of *zip-RNAi* cells; spring represents control cells which contract and pull on less elastic *zip-RNAi* clone (dashpot) during 40 min stretch. Red horizontal arrows indicate direction of stretch. * p<0.05, ****p<0.0001 with *t*-test. n.s. = non significant. Scale bars, 5 cm (A), 50 *μ*m (D), 10 *μ*m (E,G), 5 *μ*m (I).

To determine if the deformations we applied resulted in forces of a physiological magnitude, we experimentally derived the elastic index of the tissue (Young’s modulus) (Fig. S1A-E, Movie S1; see also Experimental Procedures). We subsequently calculated that the forces applied during stretching of the disc are in the range of 5-30nN per cell area and that about 30nN is required in order to produce 100% strain. This is of a similar magnitude to cellular forces previously measured *in vivo* (Stewart et al., 2011).

To test the function of MyoII in mechanical responses we generated Myosin heavy chain (Zipper) RNAi (*zip*-RNAi) clones in tissues expressing β-catenin (Armadillo) fused to GFP (Arm::GFP) (Fig. 1E, Fig. S1F). Upon application of an exogenous bidirectional anterior-posterior aligned stretch, we quantified cell shape changes in Arm::GFP-expressing tissue containing clones with *zip*-RNAi mutant cells. We used automatically-extracted changes in cell aspect ratios as a measure of cell deformation (Fig. 1E, Fig. S1G). While both the control and *zip*-RNAi tissue increased their aspect ratio significantly, indicating the tissue is stretched, the *zip*-RNAi cells became nearly two fold more elongated (Fig. 1E-F). These results imply that cells depleted of MyoII are more deformable and our aspect ratio measurements suggest that they are ~50% softer than control cells, consistent with AFM measurements on isolated cells (Martens and Radmacher, 2008). Furthermore, 10 minutes following tissue relaxation, control cells returned to their pre-stretch shape while *zip*-RNAi cells still remained significantly deformed, implying reduced elasticity of these cells (Fig. 1G-H). We could reproduce these results also by depletion of MyoII regulatory light chain, using *spaghetti squash* RNAi (*sqh-* RNAi) (Fig. S1I,J). This is in agreement with reduced Sqh levels in *zip*-RNAi cells (Fig. S1K).

To gain further insight into the physical properties of *zip*-RNAi clones we applied a defined static stretch for a longer period of time (40 min). While control cells around the clone remained at the same level of deformation throughout the timecourse, *zip*-RNAi cells continued to deform along the line of stretch (Fig. 1I-J, Movie S2). To test the prediction that the expansion of softer *zip*-RNAi clone is mediated by contraction of neighboring control cells, we next investigated the behavior of cells directly adjacent to both sides of the *zip*-RNAi clone and aligned in series with the direction of the stretch (Fig. S1H). We observed a gradual drop over time (10 min compared to 40 min stretch) in the aspect ratio change of this specific cell population, which coincided with increased elongation of *zip*-RNAi cells (Fig. S1H). The observed behavior is reminiscent of two springs (control cells) in series with a dashpot (*zip*-RNAi clone) in between them (Fig. 1K). The gradual expansion of the *zip*-RNAi population is thus a direct consequence of the control cells elastically contracting when stretched and pulling on the less elastic *zip*-RNAi cells, the viscous properties of which now dominate and dictate the gradual expansion. Taken together, these findings demonstrate that MyoII is essential for setting both the elasticity and stiffness of the wing disc epithelial tissue.

### Total and activated MyoII polarize with mechanical stretching

To gain mechanistic insight into how MyoII might govern tissue elasticity and stiffness, we looked at the localization of MyoII regulatory light chain (Sqh), Sqh::mCherry, co-expressed with a marker of cell junctions E-cadherin::GFP (E-cad) in anchored (non-stretched, just clamped in the device) and stretched discs. Strikingly, we observed that within a few minutes of stretch, MyoII became very strongly enriched on the junctions parallel to the axis of the stretch, while no change of localization was detected for E-cad (Fig. 2A-A’, Fig. S2A-A’). This phenomenon occurred irrespective of different genetic backgrounds, fluorescent tags or stretch orientation (Fig. S2B-C’). Quantification of mean junctional intensities of MyoII and E-cad (explained in Fig. S3 and Experimental Procedures) in anchored and stretched wing discs demonstrated that there was a significant enrichment of MyoII, but not E-cad on stretched (horizontal) junctions relative to unstretched (vertical) junctions, a property hereafter referred to as MyoII polarity (Fig. 2A,C). Furthermore, the relative enrichment of MyoII resulted from both MyoII increase on stretched junctions and concomitant decrease on unstretched junctions, while total junctional MyoII amount remained unchanged (Fig. 2D-F). This suggests that stretch either redistributes MyoII from unstretched to stretched junctions or changes MyoII stability on the respective junctions (Fig. 2B).

**Figure 2:**
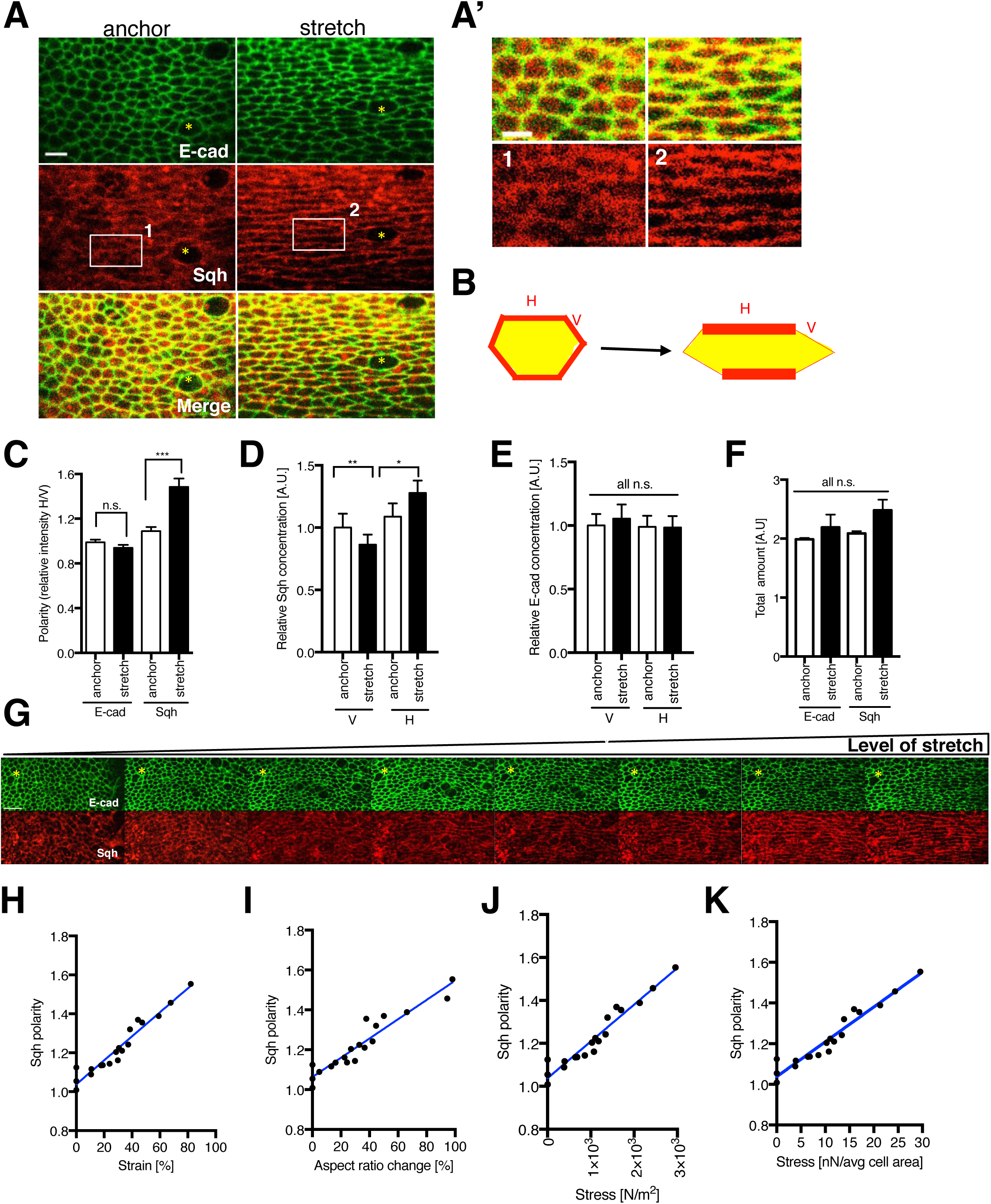
Myosin polarizes with mechanical stretch. (A) Wing disc expressing E-cad::GFP and Sqh::mCherry prior to stretch (anchor) and after stretch (15 min). (A’), Insets of discs shown in (A). (B) Schematics demonstrating change in Sqh concentration before and after stretch on vertical (V) and horizontal (H) junctions (C) Quantification of mean fluorescent intensity of E-cad::GFP and Sqh::mCherry on horizontal (H) junctions relative to the intensity on vertical (V) junctions (referred to as “polarity”) prior to and after 15 min stretch. The detailed rationale of intensity determination is described in Fig. S3 and Experimental Procedures. (D-E), Quantification of E-cad::GFP (D) and Sqh::mCherry (E) concentration (i.e. mean intensity per junctional unit area) prior and after stretch on both horizontal and vertical junctions normalized to starting concentration on unstretched (anchored) vertical junctions. (F) Quantification of total amount of Sqh::mCherry and E-cad::GFP fluorescence on vertical and horizontal junctions in anchored and stretched discs. (G) Tissue expressing E-cad::GFP and Sqh::mCherry subjected to increasing levels of stretch. (H) Quantification of Sqh::mCherry polarity (as in c) of experiment described in (G). The level of stretch is measured as a change in tissue strain (deformation relative to unstretched tissue). (I-K) Sqh::mCherry polarity emergence as a function of cell aspect ratio change (I), stress (J), stress applied on average cell area of 10 *μ*m^2^ (K). Yellow asterisks indicate equivalent regions in the tissue. N=8 wing discs (A-F), N=3 wing discs (G-K); all experiments are plotted as mean ± S.E.M., * p<0.05 with *t*-test. Scale bars, 5 *μ*m (A), 3 *μ*m (A’), 10 *μ*m (G).

To understand how the extent and duration of mechanical stretch affects MyoII polarity, we stretched the wing disc for short (up to 20 minutes) and long (up to 3 hours) periods, as well as varied the magnitude of stretch. In short time scales, we quantified that MyoII polarizes proportionally with cell deformation change, stress and strain (Fig. 2G-K, Movie S3 and S4). We also observed that both the cell shape change and polarity increase seen upon stretch were fully reversible after we relaxed the tissue (Fig. 3A-B). Interestingly, during a prolonged (up to three hour) stretch, we observed a gradual decrease in MyoII polarity, while constant cell deformation was maintained. Notably, at the end of the assay, when stretch was alleviated, the cell shape still remained 25% significantly more elongated compared to the pre-stretch aspect ratio (Fig. 3C-D), while at the same time MyoII polarity was completely lost. These findings demonstrate that MyoII polarity follows the patterns of cell and tissue deformation on short time scales. Conversely, during prolonged stretch, the uncoupling of cell shape and MyoII polarity suggests that tissue remodeling and stress dissipation allows tissue to acquire a new, preferred homeostatic shape.

**Figure 3:**
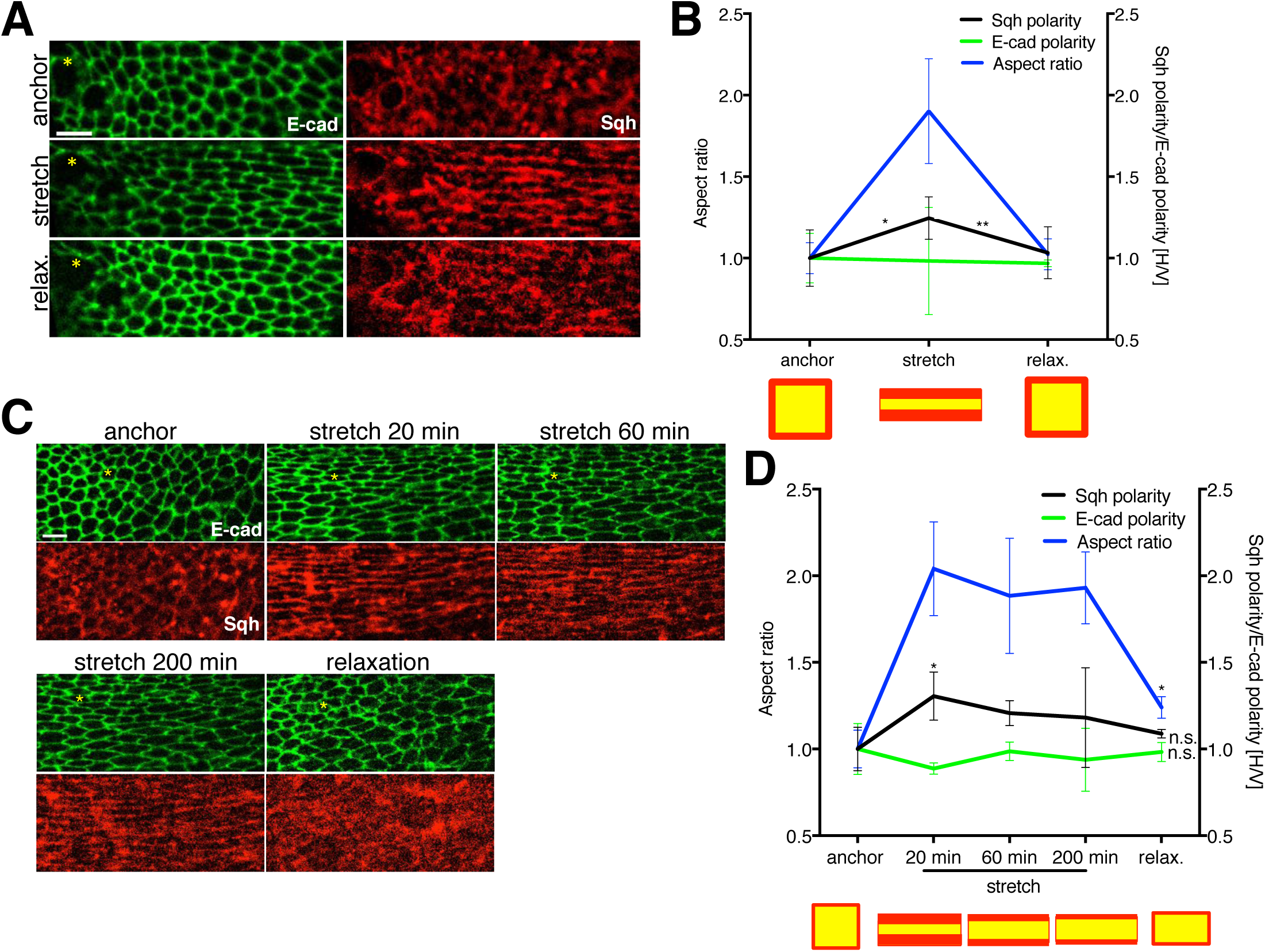
Uncoupling of tissue shape and MyoII polarity after prolonged stretch. (A) Short (15 min) stretch and relaxation (10 min) of E-cad::GFP and Sqh::mCherry expressing discs. (B) Quantification of Sqh::mCherry polarity, E-cad::GFP polarity and aspect ratio change of experiment described in (A); schematics below the graph illustrates quantified changes in cell aspect ratio (yellow) and Sqh polarity (red). Statistical test compares Sqh polarity between anchored and stretched or stretched and relaxed disc. (C) Long (200 min) stretch and relaxation (10 min) of E-cad::GFP and Sqh::mCherry expressing discs. (D) Quantification of Sqh::mCherry polarity, E-cad::GFP polarity and aspect ratio change of experiment described in (C); schematics below the graph illustrates quantified changes in aspect ratio (yellow) and Sqh polarity (red). Statistical test compares changes with respect to initial (anchored) state. Yellow asterisks indicate equivalent regions in the tissue. N=3 wing discs; all experiments are plotted as mean ± S.E.M., * p<0.05 with t-test. Scale bars, 5 *μ*m.

Importantly, as the requirement for active contractile MyoII filament formation requires prior phosphorylation of MyoII, we looked into how the activated form of MyoII behaves in mechanically perturbed tissues. To circumvent the technical problem of staining in live stretched tissues, we exploited the fact that during compression, wing discs behave elastically and stretches in the direction perpendicular to compression, concomitantly forming polarized MyoII cables in stretched cells that are under increased tension, as evidenced by laser ablation experiments (Fig. 4A-C). Such tissues can be snap fixed in the microchannels and stained while preserving its deformed shape (see Experimental Procedures). Accordingly, following compression, we could observe phospho-MyoII associated with polarized, cable-like structures (Fig. 4D-D’). Taken together, both total MyoII and activated phospho-MyoII polarize with mechanical stretch.

**Figure 4:**
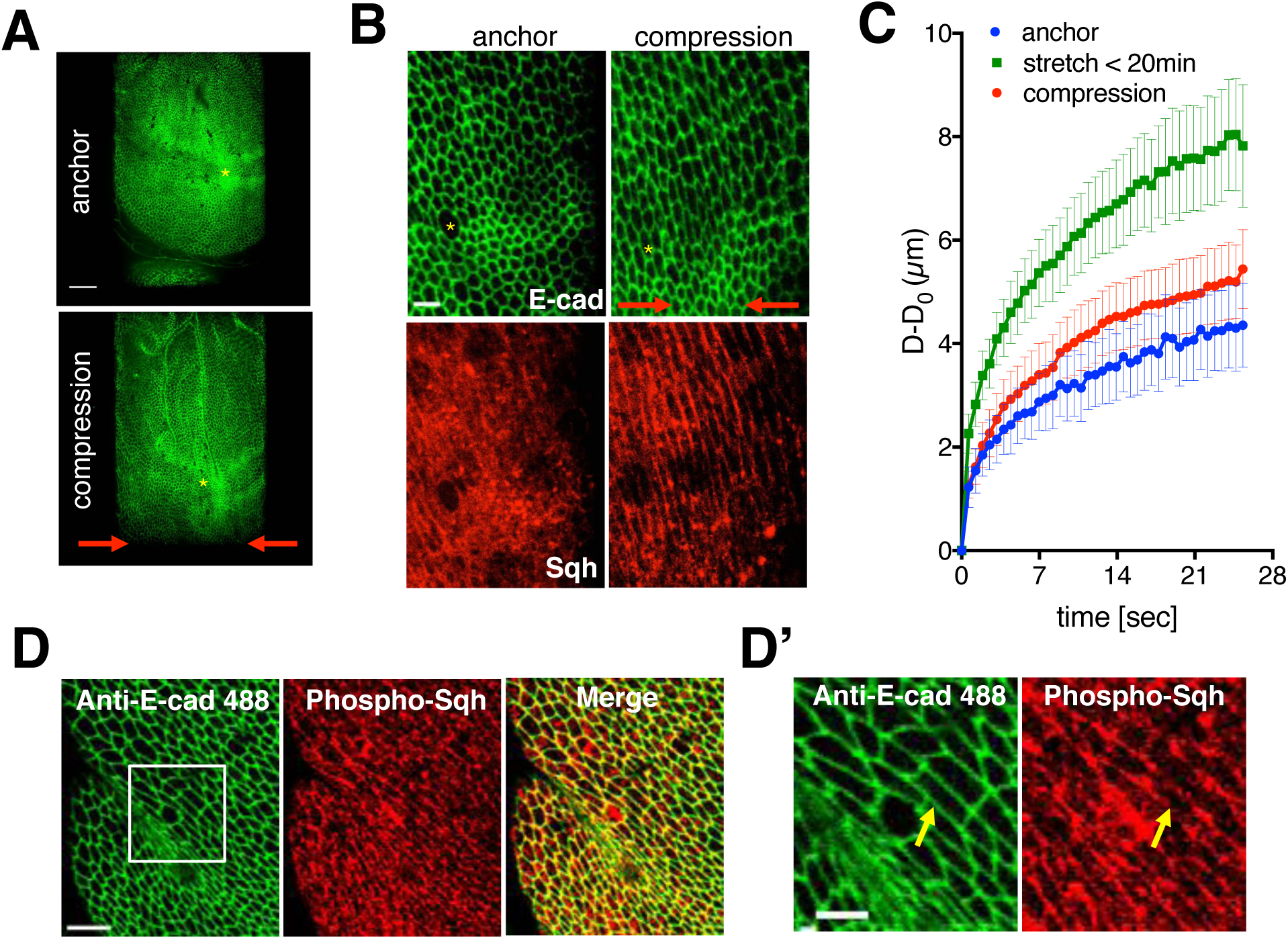
Phosphorylated MyoII polarizes in mechanically stretched cells. (A) Overview of the E-cad::GFP disc prior to (anchor) and after compression. (B) E-cad::GFP and Sqh::mCherry localization in anchored and compressed wing disc. (C) Laser cuts (perpendicular to the line of stretch) across circa 10 cells in E-cad::GFP expressing wing disc tissue subjected to anchoring, short (<20 min) stretching and short (<20 min) compression. Plot shows increase in distance between recoiling short axis [μm] of the fitted ellipse (see Fig. S5A for details) against time [sec.]; data is plotted as mean ± S.E.M, n= 7–10 discs/condition. (D) Activated Myosin (anti-phosphorylated-Sqh; in red) and anti-E-cad (green) staining in compressed discs. (D’) Close up of compression experiment described in (D); yellow arrow points towards an example of polarized phosphorylated-Sqh cable in compressed discs. Scale bars, 5*μ*m (B, D’), 10*μ*m (D), 30*μ*m (A).

### Tension increases immediately after stretch but drops gradually as the tissue remodels

To test if MyoII polarity correlates with the tension in the stretched tissue we used laser ablation to analyse both cell-level and tissue-scale forces. Using long (spanning circa 10 cells) and single junctional cuts perpendicular to the stretch, we determined that, along the direction of tissue deformation, tissue-scale and cell-scale tension increases significantly directly after stretch and decreases to the pre-stretch (anchor) levels two hours later (Fig. 5A-D, Fig. S4A-D, Movie S5 and S6). The observed drop of tension in prolonged stretch is likely a consequence of the cellular remodeling events that drive tissue relaxation, such as actomyosin remodelling, cell division and cell intercalation. Further retraction of the tissue such that cells return to their original pre-stretch shape thus effectively compresses the tissue, as evidenced by further drop in tension (Fig. 5A-B). In contrast, tissue relaxation following short (<20 min) stretch has the same tension as the anchored disc (Fig. S4C). Changing the orientation of the laser cuts to the orthogonal direction revealed that the tension was isotropic in the anchored tissue. However these junctions had significantly lower tension in the stretched tissue, indicating that the majority of tissue tension was aligned with the direction of stretch (Fig. 3E-F, Fig. S4B,E), and that the orthogonal direction experiences more compression than in the anchored state. Importantly, we could confirm that the pattern of junctional recoil is consistent with the changes in MyoII accumulation (Fig. 5D compare with Fig. 3C-D). Taken together, we can conclude that MyoII polarity correlates well with tension changes in the tissue, rather than cell shape changes.

**Figure 5:**
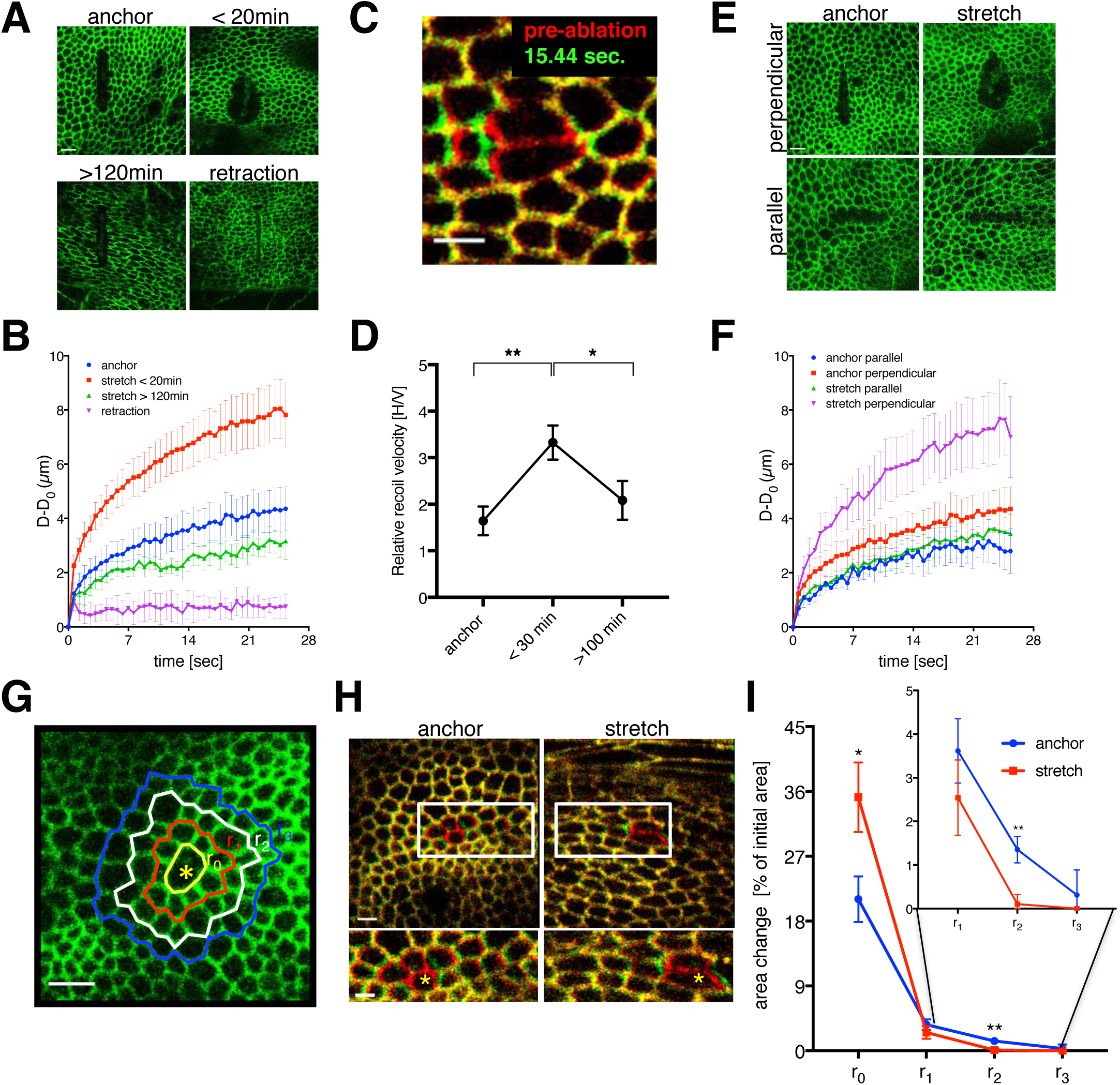
Emergent tension patterns reveal tissue stiffening upon mechanical stretch. (A) Laser cuts (perpendicular to the line of stretch) across circa 10 cells in E-cad::GFP expressing wing disc tissue subjected to anchoring, short (<20 min) and long (>120 min) stretching and stretch retraction (tissue forced back to the original, pre-stretch shape. (B) Quantification of the experiment described in (A). Plot shows increase in distance between recoiling short axis [μm] of the fitted ellipse (see Fig S4A-B for details) against time [sec.]; data is plotted as mean ± S.E.M; anchor and short stretch data plotted here is the same as in Fig. 4B, n=3–11 wing discs per condition. (C) An overlay of a junction prior to (red) and 15.44 sec. after ablation (green). (D) Plot showing relative initial recoil velocity (velocity on horizontal, stretched junctions normalized to velocity on vertical, unstretched junctions) for individual junctions ablated in anchored or stretched (<30 min, >100 min) E-cad::GFP expressing discs. Data is plotted as a mean ± S.E.M, n=17–24 cuts per condition. (E) Laser cuts (perpendicular or parallel to the line of stretch) across circa 10 cells in E-cad::GFP expressing wing disc tissue subjected to anchoring or short stretching (< 30 min).. (F) Quantification of experiment described in (E). Data is quantified and presented as in (B). N=4-7 wing discs per condition. (G) Image depicting injury site (yellow asterisk) and selection of cell rows used for tissue recoil propagation measurement; r: row. (H) An overlay of anchored and stretched discs prior to ablation (red) and 15.44 seconds post ablation (green). Inset zooms in on the area next to ablation site (marked with yellow asterisk). (I) Recoil area (% increase from pre-ablation area) for row 0, row 1, row 2 and row 3 in stretched and anchored discs. Data is plotted as mean ± S.E.M, n=8-13 discs/condition. Error bars indicate S.E.M.; * p<0.05 and **p<0.01 with *t*-test. Scale bars, 5 *μ*m (A,E,G), 3 *μ*m (C,H), 2 *μ*m (H: inset).

### Stretch-induced MyoII cables rigidify the tissue

Junctional laser ablations not only reveal the tensile state of tissues but also lead to micro-injuries in the cells. We wondered whether we could observe any differences in tissue responses to injury in anchored and stretched tissue (see Experimental Procedures). Interestingly, we noticed two types of behaviors depending on the proximity to the injury site. Locally, at the level of injured cells (row 0) we noticed significantly higher tissue recoil in stretched cells, consistent with the measured increase in tension (Fig. 5G-I, Fig S4B). Remarkably, in cell rows further away from the injury site (row 2 and row 3) this relationship became reversed, with anchored cells demonstrating higher degree of recoil compared to stretched tissue (~100% more recoil in rows 2 and 3) (Fig. 5G-I). This is despite much higher global tension in the latter condition (Fig. S4A-C). It thus appears that tensile stress stiffens the tissue to buffer mechanical perturbations such that they fail to propagate beyond the site of injury. Given that MyoII polarization contributes to the stiffness of the tissue (Fig. 1E-F), it is likely that this stiff tissue lattice is driven by stretch-induced tissue scale MyoII polarization. Together, our data strongly suggest that the presence of polarized MyoII cables stiffens the tissue to preserve tissue morphology and to prevent local tissue damage from propagating.

### MyoII polarity is Rho-Kinase independent

Previous work has shown that assembly of contractile MyoII filaments is regulated by direct phosphorylation of myosin light chain by Rho-Kinase (Rok), through the Rho1 pathway (Riento and Ridley, 2003). To uncover the molecular mechanism driving stretch-induced MyoII polarity, we therefore examined Rok function in wing disc mechanical responses. Surprisingly, we observed that while both Rho1::GFP and the Rok reporter, Rok^K116a^::Venus became polarized with stretch, the loss of Rok *(enGal4>rok RNAi* and *Drok*^2^ loss of function clones) did not prevent MyoII polarization (Fig. 6A-C, Fig. S5E). For both *rok* RNAi and *Drok*^2^ conditions, we observed an increase in cell size, a decrease in overall MyoII levels, and a decrease in cell tension as evidenced by laser junction ablations, indicative of successful Rok inactivation (Fig. S5A-D, Movie S7 and S8). To investigate whether observed Rok polarity is in fact a consequence of MyoII polarization we generated *zip-RNAi* clones in tissue expressing Rok^K116a^::Venus and subjected it to stretch. Remarkably, induction of *zip-RNAi* clones completely abolished polarization of Rok (Fig. 6D). We therefore conclude that stretch-induced MyoII polarity is Rok independent, while Rok polarity emerges from MyoII polarity.

**Figure 6:**
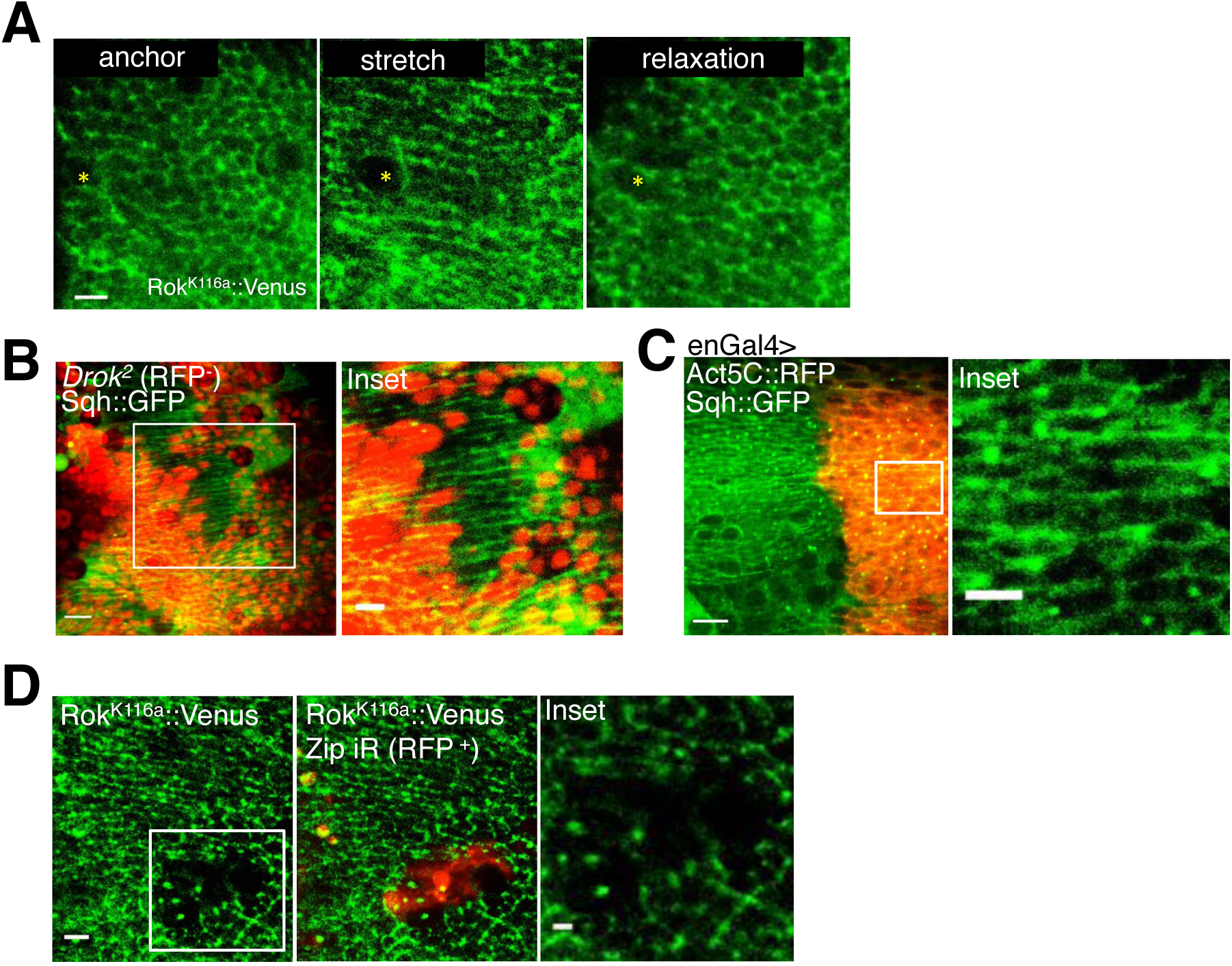
Myosin polarity is Rok independent. (A) Rok^K116a^::Venus localization in anchored, stretched, relaxed third instar imaginal discs. (B) *Drok^2^* mitotic clones (RFP^-^) in Sqh::GFP third instar imaginal disc. Inset demonstrates Sqh::GFP polarity inside *Drok^2^* clone. (C) *Sqh::GFP, enGal4>UAS-rok RNAi, UAS-Act5CRFP* discs subjected to stretching; inset demonstrates Sqh::GFP polarity in *rok* RNAi cells (marked by red. (D) *zip* RNAi clones (labeled as zip iR, indicated in red) in stretched third instar Rok^K116a^::Venus discs. Error bars indicate S.E.M.; * p<0.05 and **p<0.01 with *t*-test. Scale bars, 2 *μ*m (D: inset), 5 *μ*m (A,D, B: inset, C: inset), 10 *μ*m (B,C).

### MyoII polarization requires the dynamic remodeling of linear actin filaments generated by formins

To identify the upstream regulator of stress-induced MyoII polarity, we performed an RNAi and gene overexpression (OE) screen, targeting a selection of potential or reported MyoII regulators (Fig. S6A and Experimental Procedures). While Sqh::GFP was expressed ubiquitously, the *engrailed-Gal4 (en-Gal4)* driver targeted RNAis/OE to the posterior side, marked by expression of *UAS-Actin5C::RFP* (Act5C::RFP). We confirmed that Act5C::RFP on its own had no visible effect on MyoII polarity (Fig. S6B-B’). A positive result was defined as a failure to polarize Sqh::GFP upon stretch, while retaining junctional Sqh (Fig. S6A; see Experimental Procedures for details). Unexpectedly, none of the reported Myosin regulators such as Myosin Light Chain Kinase (MLCK) were required for stretch-induced MyoII polarity. Instead, we observed polarization defects when we depleted the formin Diaphanous, an actin nucleator that generates linear arrays of F-actin, which form scaffolds where contractile MyoII can be assembled (two different RNAi lines, Fig. 7A-A’, Fig. S6C-C’,F) (Jegou et al., 2013). This polarity defect was further quantified in discs co-expressing E-cad::GFP (Fig. 7B-D). To rule out unspecific effects resulting from an increased size of *dia*-RNAi cells, we confirmed that depletion of Pebble and Rok, which have cytokinetic and cell-size defects similar to those seen in Dia-deficiency, did not produce any MyoII polarity defects (Fig. 6B-C, Fig. S5A-C, Fig. S6F). Depletion of F-actin depolymerisation factors cofilin and AIP1 also resulted in MyoII polarization defects (Fig. 7E-E’), suggesting that existing F-actin filaments must also disassemble for force-induced MyoII polarization (Chen et al., 2015; Nadkarni and Brieher, 2014). Consistent with this, we observed that the actin reporters, Utrophin::GFP and Act5C::RFP became polarized and Act5C::RFP co-localized with MyoII cables upon application of the mechanical force (Fig. S6G-H). Finally perturbation of branched F-actin, which is believed to be a less favourable scaffold for myosin contractility, by depletion of Arp2, had no effect on MyoII polarity (Fig. S6F). Taken together our data strongly suggest that the dynamic remodelling of linear actin cables generated by formins is required for MyoII polarization upon stretch. This is in agreement with previous *in vitro* studies that demonstrated that Diaphanous is tension sensitive whereas Arp2/3 is not (Jegou et al., 2013; Lomakin et al., 2015).

**Figure 7:**
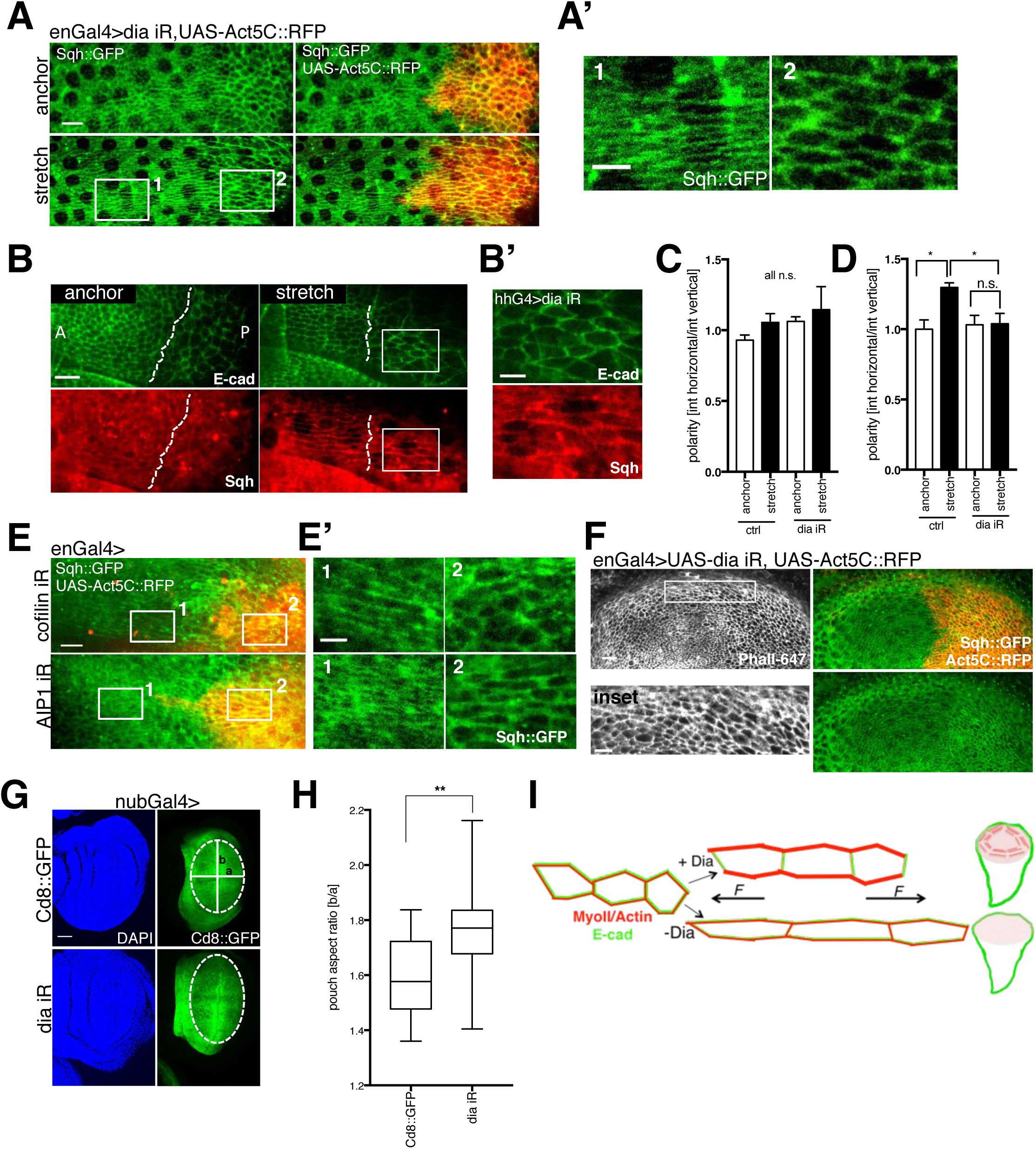
MyoII polarity is required for wing disc shape maintenance via Dia-dependent actin polymerization. (A) Discs expressing Sqh::GFP and *enGal4>UAS-dia RNAi, UAS-Act5c-RFP* that were stretched; UAS-Act5C::RFP marks *dia* iR (posterior side) in red. (A’) Insets comparing Sqh::GFP polarity in stretched *enGal4>dia iR, UAS-Act5c-RFP* discs; panel 1 refers to anterior (ctrl) side; panel 2 refers to posterior *(dia* iR) side. (B) Tissue marked with E-cad::GFP and Sqh::mCherry, expressing *dia-RNAi (dia* iR) with hhGal4 driver and subjected to stretching. Dotted line indicates anterior-posterior (A-P) compartment boundary. (B’) Inset demonstrates lack of Sqh::GFP polarity in *dia-RNAi* cells. (C-D), Quantification of E-cad::GFP polarity (C) and Sqh::mCherry polarity (D) in experiment described in (B); polarity is measured as mean fluorescent intensity on horizontal junctions relative to vertical junctions (see Experimental Procedures for details); n=4 wing discs. (E) Stretched discs expressing Sqh::GFP and *enGal4>UAS-cofilin* (upper panel) *or UAS-AIP1* (lower panel) *RNAi* and *UAS-Act5c-RFP* (marks posterior side in red). (E’), Insets comparing Sqh::GFP polarity in stretched discs described in (E); panel 1 refers to anterior (ctrl) side; panel 2 refers to posterior *(dia* iR) side. (F) Sqh::GFP polarity in enGal4> *dia* iR, UAS-Act5C::RFP discs labeled with Phalloidin-647; Inset (Phall-647) zooms in on cell shape in anterior and posterior disc compartments. (G) Discs expressing nubGal4 and Cd8::GFP (control) or *dia* iR (KK) labeled with DAPI; ellipse is fitted into the pouch region (red dotted line) and aspect ratio determined as long ellipse axis (b) divided by short ellipse axis (a) (white lines). (H) Box plot showing distribution of wing disc pouch aspect ratios in conditions described in (G), n=19 (ctrl) and 16 (dia iR) wing pouches; horizontal line indicates median. (I) Model of Diaphanous-Myosin force buffering pathway. Tissue (E-cad marks cell junctions in green) subjected to mechanical stretching polarizes MyoII and Actin (in red) in Dia-dependent manner. This mechanism is required to limit tension induced cell deformation and thus maintain wing disc pouch shape. Error bars indicate S.E.M.; * p<0.05 and **p<0.01 with *t*-test. Scale bars, 5 *μ*m (A’,B’,E’, F: inset), 10 *μ*m (A,B,E), 30 *μ*m (F, G).

### MyoII polarity pathway is essential to maintain tissue shape

Having identified a condition that specifically decreases MyoII polarity we next tested its impact on cell deformation upon mechanical stretch. In agreement with our MyoII RNAi data (Fig. 1E-F), Diaphanous depletion significantly increased cell deformation (Fig. S6D-E). Taken together, our results showed that Diaphanous is required for induction of MyoII polarity and subsequent cell shape maintenance in tissues challenged with extrinsic mechanical stress.

To test the requirements of Dia-dependent MyoII polarization in the context of mechanical forces emerging *in vivo*, we explored previous reports showing that mechanical tension emerges in proximal wing disc cells as a result of proliferation anisotropy (Legoff et al., 2013; Mao et al., 2013). Interestingly, the increased tension was shown to correlate with polarization of phosphorylated MyoII in these cells (Legoff et al., 2013; Mao et al., 2013). However, whether MyoII polarized as a direct response to mechanical force was never shown. Our data showing that MyoII polarizes as a direct response to exogenous stretch suggests that the same mechanism is occurring during *in vivo* wing disc development. To test a causative link between proliferation induced tension and MyoII polarity we induced ectopic overgrowth by generating highly-proliferative clones mutant for *warts (wts^x1^)* in the wing disc notum, where no prior emergent patterns of MyoII polarity could be detected (Fig. S7A). Following 72h induction of clone growth, we could observe concomitant cell stretching and Sqh::GFP polarized cables forming around the overgrowing *warts* mutant clone (Fig. S7B-B’). We could thus confirm that proliferation induced stretching forces are sufficient to promote MyoII polarity *in vivo*. This result is in agreement with the previous observation by LeGoff *et al*., showing that induction of overgrowth by overexpression of the Hippo pathway targets, Yorkie or Expanded was correlated with MyoII anisotropy in the wing disc (Legoff et al., 2013).

Having established a causal link between emergent tension patterns in the wing disc and MyoII polarity, we next examined the function of Dia in this process. To this end we targeted *dia*-RNAi expression to the posterior compartment of the disc. In agreement with exogenous stretching data, we observed defective Sqh::GFP polarization of the proximal cells in the posterior compartment when compared to proximal cells in anterior (control) compartment (Fig. 7F) (Mao et al., 2013). At the same time cell stretching was at least as high in *dia-*RNAi as in control cells as evidenced by Phalloidin staining (Fig. 7F: inset, Fig. S7C-C’). This result indicated that MyoII polarity was compromised in *dia* depleted cells *in vivo*.

Due to the emergence of mechanical tension in developmental growth one would predict that the lack of MyoII polarization mechanism could impact on tissue shape. To test this prediction we targeted *dia-*RNAi expression to the whole wing disc pouch with *nubbin-Gal4* driver (nubGal4). Comparing control and *dia-*RNAi discs we could not observe any significant difference in the overall size (Fig. S7D). Strikingly though, *dia-*RNAi wing disc pouches were significantly wider as measured by increase in their aspect ratio, indicative of increased tissue deformation (Fig. 7G-H). In summary, the physiological role of Dia-MyoII pathway is to limit tension-induced cell deformation by inducing MyoII polarity (summarized in Fig. 7I).

## Discussion

Our study has uncovered a novel MyoII polarity pathway that allows fast and efficient tissue-scale adaptation to mechanical perturbations. We demonstrated that mechanical stretch induces MyoII polarization, a process that has a dual role in stretch responses by i) maintaining tissue elasticity and ii) setting tissue stiffness to protect against fractures and injuries. Both of the identified functions ensure that tissue shape and subsequent function are preserved while mechanical stress is being dissipated. Importantly, the core of the pathway, formation of polarized MyoII cables, does not follow the canonical rules of MyoII upstream biochemical activation via Rho1-Rok, seen in processes like embryonic junctional remodeling, but rather stems from more fundamental, mechanosensitive properties of actomyosin complexes (Bertet et al., 2004; Rauzi et al., 2010). Prior *in vitro* studies have demonstrated that application of mechanical tension to isolated actin filaments increases the rate of their polymerization, a phenomenon, dependent in part on the barbed-end associated actin nucleator Diaphanous (Courtemanche et al., 2013; Galkin et al., 2012; Jegou et al., 2013; Kovacs et al., 2007; Kozlov and Bershadsky, 2004). Such tensed actin filaments could also stabilize more ADP-bound Myosin (Llinas et al., 2015). Whether this mechanism is sufficient to polarize MyoII *in vivo* is unclear. If this mechanism operates *in vivo*, extrinsic forces applied on the tissue would stimulate actin polymerization via formins in the direction of stretch. These new filaments would recruit more Myosin, which could have a scaffolding role for Rok and Rho1. The observed polarization of Rok and Rho1 could serve as an emergent positive feedback signal to further enhance or stabilize polarized actomyosin complexes rather than being the instructive upstream cue.

Gradual changes in mechanical properties such as tissue stiffness are a normal part of development as exemplified in the differentiation of embryonic stem cells (Butcher et al., 2009). Here we show how epithelial tissues can rapidly refine their mechanical properties as an adaptive and protective response to the constant fluctuations in physical forces from a tissue’s environment. This response buffers mechanical stresses to maintain the balance of intrinsic tissue forces and preserve tissue integrity. This fundamental mechanosensitive pathway could play a broad role in force adaptation in both development and pathological situations such as cancer and tissue repair, where tissue mechanics are affected in an uncontrolled manner.

## Experimental Procedures

### Fabrication of Silicon Master Mould

Optical lithography was performed on silicon wafers and carried out using the cleanroom facilities of the London Centre for Nanotechnology. Silicon substrates were cleaned through repeated washes with acetone and isopropanol, and covered with Su-8 2050 (Microchem). The photoresist was spun with a spin-coater, according to the manufacturer’s instructions, to achieve a uniform thickness of 50*μ*m. The soft-bake step, required after spin-coating to evaporate the excess of solvent and to densify the resist, was performed by placing the wafers for 3 minutes on a hot-plate at 65°C and, sequentially, at 95 °C for 8 minutes. The photoresist is exposed to a pattern of intense UV light through a photomask. The UV exposure time needed to crosslink the SU-8 was calculated by dividing the exposure energy (mJ/cm^2^) indicated in the Su-8 data sheet by the light intensity of the Karl Suss MJB3 mask aligner (20 mJ/cm^2^). Post-exposure bake was performed on hot-plates, at 65°C (2 minutes) and at 95°C (8 minutes). The wafer was placed in Microposit™ EC solvent for development for approximately 6 minutes, and finally hard-baked at 180°C for 5 minutes, to stabilize and harden the developed photoresist.

### Preparation of patterned polydimethylsiloxane (PDMS) membranes

PDMS mix was prepared according to the manufacturer’s recommendations (SYLGARD 184 elastomer kit, Dow Corning), briefly spinned down to remove bubbles (2min, 1000rpm) and let to cure at room temperature for 4h prior to spinning (30 seconds, 900 rpm) on a patterned silicon wafer mould (microchannels of 80-120*μ*m width and 50*μ*m depth) with Spin Coater (SPS Spin 150). The PDMS covered mould was then heated up to 100° C for 10 min to allow full PDMS curing and the PDMS was subsequently peeled off the wafer and transferred onto a transparency sheet for storage.

### Stretcher design

A custom made stretching/compression device was constructed. In detail, the basic drive for the unit consists of a combination of linear bearings with left and right-handed fine pitch screw threads to provide the desired movement. The extended metal arms connect the drive to the clamping mechanism. Moving the arms apart (inside-out) or bringing them closer together (outside-in) respectively stretches or compresses the clamped material. The device fits onto most standard microscope stage plate holders suitable for imaging.

For stretching/compression assays, a pre-patterned PDMS membrane (Fig. 1b) was spread over two metal arms, overlaid with a plain (unpatterned) PDMS membrane (GelPak, 6.5 mil) and clamped on both ends. An isolated wing disc was suspended in the media, injected in between two PDMS layers and positioned with forceps over a microchannel, with its apical side of wing disc proper facing down the channel. A PDMS chamber was placed on top of the membranes and filled with media to prevent the drying out of the sample.

### Stretching experiments

If not specified otherwise stretching was performed bi-directionally along anterior-posterior (A-P) axis. An anchored disc was defined as a disc injected in between PDMS membranes but not subjected to any mechanical perturbation. Short stretch was defined as less than 30 minutes and long stretch as more than 120 min. Relaxation was achieved by bringing the stretcher arms back to the pre-stretch position. For assessment of degree of polarity with respect to applied strain (Fig. 2) discs were stretched to increasing degrees with 5 min pause intervals in between to allow for the full emergence of MyoII polarization. We confirmed stretching had no effect on discs viability as cells continued dividing at least 1 hour post-stretch (see Movie S3). For compression (Fig. 4), PDMS membranes were pre-stretched prior to disc injection. Disc was subsequently dropped inside the channel and compressed by bringing the stretcher arms back. In cases where immunofluorescence staining was required (only applicable to compression experiments), the upper PDMS layer was omitted and instead the discs were directly fixed on the device with 18% PFA.

### *Drosophila* strains and genetics

Flies of the following genotype were used: *Ecad::GFP* (knock-in(Huang et al., 2009)); *pnrGal4, Sqh::mCherry* (gift from Baum lab), *sqhP-Utrophin::GFP(Rauzi et al., 2010)*, *Sqh::GFP, enGal4; UAS-Act5C::RFP* (self-generated), *Rho1::GFP* (protein trap, Kyoto Stock Centre, DGRC ID: 110833), *SqhAx3;; Sqh::GFP* (Jordan and Karess, 1997), *Rok^K116a^::Venus* (Simoes Sde et al., 2010), *ubi-Ecad::GFP; hhGal4, UAS-IAP/TM6* (gift from N. Tapon), *Sqh::mCherry* (III)(Martin et al., 2009), *UAS-Cd8::GFP(Mao et al., 2013), nubGal4(Mao et al., 2013), UAS-torso^D^/βcyt (Myospheroid mutant)(Martin-Bermudo and Brown, 1999), ubi-mRFP.nls,w,hsF,Frt19A* (Bloomington, 31418), *yw,Drok^2^,Frt19A/FM7c* (Bloomington, 6666)(Escudero et al., 2007), *ubi-mRFP.nls,w,hs.flp,Frt19A;; Arm-GFP* (self-generated), *ubi-mRFP.nls,w,hs.flp,Frt19A;; Sqh-GFP* (self-generated), *yw hs.flp; E-cadherin::GFP; FRT82B ubi-mRFP-nls* (Mao et al., 2013), *yw;; FRT82B wtsX1/TM6b* (gift from Tian Xu)(Mao et al., 2013), *actin-FRT-stop-FRT-Gal4-UAS-Cd8::mCherry; Arm::GFP* (self-generated), *actin-FRT-stop-FRT-Gal4-UAS-Cd8::mCherry; Sqh::GFP* (self-generated), *actin-FRT-stop-FRT-Gal4-UAS-Cd8::mCherry; Rok^K116a^::Venus* (self-generated), *yw hs.flp; UAS-zip iR/SM6-TM6* (self-generated).

### Clones generation

For *Drok^2^* mitotic clones *ubi-mRFP.nls,w, hs.flp,Frt19A/ yw,Drok^2^,Frt19A; Arm-GFP/+* or *ubi-mRFP.nls,w,hsF,Frt19A/ yw,Drok^2^,Frt19A; Sqh-GFP/+* larvae were heat shocked at 37 °C for 1h at the age 48h AEL and discs were dissected for experiments 72h later (at 120h AEL). For *wts^X1^* mitotic clones yw *hs.flp; Sqh-GFP/+; FRT82B wtsX1/ FRT82B Ubi-mRFP-nls* larvae were heat shocked at 37 °C for 1h at the age 48h AEL and discs were dissected for experiments 72h later (at 120h AEL). For zipper iR flipout clones *yw hs.flp/+; actin-FRT-stop-FRT-Gal4-UAS-Cd8::mCherry /zipiR; arm-GFP/+* or *yw hs.flp/+; actin-FRT-stop-FRT-Gal4-UAS-Cd8::mCherry /zip iR; Sqh-GFP/+, yw hs.flp/+; actin-FRT-stop-FRT-Gal4-UAS-Cd8::mCherry/zip iR; Rok^K116a^::Venus/+* larvae were heat shocked at 37 °C for 13 min at the age 62–72h AEL and discs dissected for experiments 48h later (at 110-120h AEL).

### RNAi lines

All RNAi lines were purchased from Vienna *Drosophila* Resource Centre (VDRC) or National Institute of Genetics (NIG). The RNAi lines were used to silence following genes: *zipper* (VDRC, ID: 7819), *spaghetti squash* (NIG, ID: HMS00830 and HMS00437), *diaphanous* (VDRC, ID: 103914 KK and 20518 GD), *pebble* (gift from A. Mueller), *arp2* (VDRC, ID: 101999 and NIG, ID: 9901-R), *rok* (VDRC, 104675KK), *enabled* (VDRC, ID: 43058), *e-cadherin* (VDRC, ID: 103962), *drak* (VDRC, ID:107263), *src64B* (gift from N.Tapon), *src42A* (gift from N.Tapon), *zyxin* (NIG ID: 32018R-1 and 32018R-3), *mlck* (VDRC, ID:109937KK), *paxillin* (VDRC, ID:107789), *vinculin* (VDRC, ID: 34586), *α-actinin* (VDRC, ID:107263), *cofilin* (VDRC, ID: 100599), *AIP1* (VDRC, ID: 22851).

### Immunofluorescence

Discs were dissected in ice cold PBS and fixed for 30 min in 4% paraformaldehyde, washed 4x10 min with PBT (PBS, 0.3% Triton X-100), blocked for 1h with 0.5% BSA PBT and stained with primary (over night at 4°C) and fluorescently conjugated secondary antibodies (1h at room temperature). For phospho-Sqh staining discs were dissected in the media, compressed and fixed with 18% PFA for 10 min. Primary antibodies used were rat anti-E-cad (Developmental Studies Hybridoma Bank), rabbit anti-Diaphanous(Afshar et al., 2000), mouse anti-Engrailed (Developmental Studies Hybridoma Bank), mouse anti-Sqh (gift from R. Ward)(Zhang and Ward, 2011), rabbit anti-phospho-Sqh (3671, Cell Signalling). Dyes used were DAPI (D8417, Sigma) for DNA staining and Phalloidin Alexa-555 (A34055, Life technologies) or Phalloidin Alexa-647 (A22287, Life Technologies) for actin labelling. All secondary antibodies were from Life Technologies and Jackson ImmunoResearch.

### Young’s modulus determination

Force measurement device used for Young’s modulus determination was modified from previous design(Harris et al., 2012; Wyatt et al., 2015). In contrast to published device, the capillary was not glued to the NiTi wire, to allow for the arm to be flexible. The mechanical testing setup was modified to allow direct force measurements with force transducer rather than inferring it from the bending of the wire in the images.

A wing disc was immobilized with CellTak (Corning) over two coverslips attached to flexible and static rods and subsequently overlaid with culture media. Stretching force was applied with a motorized micromanipulator (Physik Instrumente). Initially, the wing disc was preconditioned, by stretching to 0.1mm at 0.01mm/s strain rate. This was repeated 5 times. Then the disc was rested for 2.5 min to allow full relaxation. The stretching experiments were conducted by stretching the discs to 0.3mm, 0.5mm. 0.65mm and 0.8mm at 0.01mm/s strain rate. The wire was calibrated by stretching it to 0.8mm at 0.01mm/s strain rate. This was repeated 3 times. For Young’s modulus determination the recorded force and measured cross sectional area (depth dimension averaged as 50 *μ*m; width measured in each of experiments) was used to derive stress. The Young’s modulus was further determined from the linear part of the stress-strain curve.

### Live imaging

For all live imaging experiments discs were cultured *ex vivo* as described previously(Heller et al., 2016; Zartman et al., 2013). Live imaging of stretching and compression experiments was performed on a Zeiss Spinning Disc confocal microscope equipped with Andor Zyla 4.2. PLUS sCMOS camera. The field of view for imaging was confined to the pouch and part of the hinge region of the wing disc with a 40x lens and 1 *μ*m depth resolution.

### Fixed sample imaging

Fluorescent imaging of fixed samples was performed on Leica Sp5 and Sp8 inverted confocal microscopes. All images were acquired with 40x objective and depth resolution of 1*μ*m.

### Analysis of junctional intensities and cell aspect ratio

Images were segmented and analysed with EpiTools software as reported before (Heller et al., 2016). In short, image was segmented (Epitools) after background subtraction (rolling ball radius 20, Fiji) and underlying junctional intensities (E-cad::GFP and Sqh::mCherry) were extracted with Cell Graph (EDGE_COLOR_TAG overlay) (Fig. S3). The following parameters were applied: 2 pixel edge intensity buffer around the junctions, selection mode 1 (tricellular signal removed), upper 50% (0.5) of the signal intensity. Junctions were selected manually and color-coded as horizontal (junction angle <45°) and vertical (junction angle >45°); resultant intensity data was exported. Polarity was defined as fluorescent mean intensity on horizontal (stretched or equivalent in anchored) junctions divided by intensity on vertical (unstretched or equivalent in anchored) junctions. Total MyoII and E-cad was quantified by multiplication of mean intensity by the junctional length. Concentration was defined as mean fluorescent intensity per unit junctional area. For aspect ratio the analysis was performed as shown previously using ELLIPSE_ELONGATION_RATIO or CELL_COLOR_TAG overlay in Cell Graph (Heller et al., 2016).

### Laser ablations

### Drok^2^ clones

Laser ablations were performed on a Zeiss LSM780 inverted two-photon microscope with a Chameleon Ultra II laser set to 730 nm and 100% power with ~16 *μ*s dwell time and × 10–12 digital zoom with 40x lens. Timelapse was recorded for a single channel (E-cad::GFP) with 780 ms scan time. For each of the experiments the overview picture was taken (E-cad::GFP, RFP) to identify the position of the clones prior to laser cutting; care was taken to take comparative regions in the disc for control and *DRok^2^* clones.

### Stretching experiments

Laser ablations were performed on a Zeiss LSM880 inverted multiphoton microscope with a Chameleon Ti:Sapphire laser set to 720 nm and 60% power and digital zoom 10x (single junctions ablations for Fig. 5C-D), 7x (single junction ablations for analysis in Fig. 5G-I), 5x (large cuts for tissue scale tension measurements; Fig. 5A,B,E,F). The single junction ablations were acquired with a Zeiss Airyscan detector set to a superresolution mode. Timelapse was recorded for a single E-cad::GFP channel with 633.02 ms (confocal imaging) or 643.47 ms (superresolution imaging) scan time. Single junctions ablations were analysed as described previously (Mao et al., 2013). For tissue scale tension measurements, an ellipse was fitted over the ablated region with Fiji. The change in short ellipse axis over time was taken as a measure of tissue recoil (Fig. S4A-B.). For determining tissue-scale ablation recoil (Fig. 5G-I) a single junction was ablated and timelapse was recorded for a single E-cad::GFP channel with 643.47 ms scan intervals. Based on the resultant videos the area was drawn around 0,1,2, and 3 rows of cells away from injured cells prior to ablation (0 sec.) and post ablation (15.4 sec.) (Fig. 5G). The difference in area normalized to pre-ablation area was taken as a recoil propagation measure (post ablation area – pre-ablation area /pre-ablation area). If required the images were corrected with stackreg Fiji plugin to account for unspecific drift.

### Screen design

*Sqh::GFP, enGal4> UAS-Act5C::RFP* line was generated for screening and crossed to respective RNAi or overexpression lines. Discs were imaged in anchored and stretched (15 min) configuration. A hit was defined on the basis of Sqh::GFP localization: uniform junctional when anchored on both posterior and anterior sides; polarized on anterior side (control) and uniform on posterior side (RNAi) when stretched.

### Wing disc pouch shape analysis

nubGal4>UAS-Cd8::GFP was used to define the areas of the wing disc pouch and to drive RNAi. An ellipse was fitted with Fiji selectively to the pouch region and aspect ratio (length of ellipse long axis/length of ellipse short axis) determined.

### Statistical analysis

No statistical methods were used to predetermine sample size. The minimum requirements for sample sizing were as follows: 1. Experiments performed on 3 independent days; 2. Wing discs from at least 3 independent animals/conditions; 3. On average 20-200 cells were analyzed per wing disc. Exclusion criteria: 1. Uneven and unspecific fluorescent signal across the sample; 2. Much higher variability when compared to other samples in the same condition group. Random wing discs were chosen within a given condition group.

For statistical analyses two-tailed unpaired or paired Student’s t-tests were used as indicated in the respective figure legends along with the exact n values used for each of the experiments. The following statistical significance cut off was applied: * p<0.05, **p<0.01, ***p<0.01, ****p<0.0001.

## Data availability

All representative data generated during this study is included in this published article. All data used for quantifications is available upon request.

## Author Contributions

Author contributions: MD designed the research, performed experiments, analyzed the data and wrote the article. NK designed and performed experiment described in Fig. S1A-E. NC and MP designed the prototype of the stretching/compression device. AB prepared silicon wafer moulds. GC and BB provided input into research design and data analysis. YM designed the research, designed the prototype of the stretching/compression device, analyzed the data, and wrote the article.

## Acknowledgements

We are grateful to Nic Tapon, in whose lab the development of the stretching device was initiated, for his support. We thank GREM (Griffon Gravure) for building the first prototype of stretching/compression device. We thank Duncan Farquharson, Simon Townsend, Piotr Sienkiewicz from Mechanical and Electronic Workshops at University College London for design and execution of final version of stretching/compression device. We thank Davide Heller for help with junctional myosin intensity measurements. We thank John Zhang for his help drawing the photo masks, and JDPhoto-Tools for printing. We thank all the authors and Shiladitya Banerjee, Martin Raff, Robert Tetley, Franck Pichaud, Nic Tapon, David Ish-Horowicz for revision of this manuscript. This work was supported by MRC funding to the MRC LMCB University Unit at UCL, award code MC_U12266B. MD is funded by a Marie Skłodowska-Curie Horizon 2020 Individual Fellowship: MRTGS. YM is funded by a Medical Research Council Fellowship MR/L009056/1 and a UCL Excellence Fellowship. NK is funded by the Rosetrees Trust, the UCL Graduate School, the EPSRC funded doctoral training program CoMPLEX and a UCL Overseas Research Scholarship. GC is supported by a consolidator grant from the European Research Council (MolCellTissMech, agreement 647186). AB is funded through EPSRC. BB is supported by CRUK 17343, BBSRC BB/K009001/1, BBSRC BB/J008532/1 grants. MP and NC are funded through CNRS and Institut Curie.

